# Millstone: Software for Multiplex Microbial Genome Analysis and Engineering

**DOI:** 10.1101/086868

**Authors:** Daniel B. Goodman, Gleb Kuznetsov, Marc J. Lajoie, Brian W. Ahern, Michael G. Napolitano, Kevin Y. Chen, Changping Chen, George M. Church

## Abstract

Inexpensive DNA sequencing and advances in genome editing have made computational analysis a major rate-limiting step in adaptive laboratory evolution and microbial genome engineering. We describe Millstone, a web-based platform which automates genotype comparison and visualization for projects with up to hundreds of genomic samples. To enable iterative genome engineering, Millstone allows users to design oligonucleotide libraries and create successive versions of reference genomes. Millstone is open source and easily deployable to a cloud platform, local cluster, or desktop, making it a scalable solution for any lab.

## Background

Microbial populations can harbor a staggering amount of genomic diversity, enabling them to evolve and adapt to diverse environments. Adaptive laboratory evolution uses this process to generate strains that are useful for biotechnology or for answering fundamental biological questions (Dragosits and Mattanovich 2013). In addition to harnessing natural variation, biologists can generate targeted genomic diversity in a population of cells and then screen or select for phenotypes of interest (Wang et al. 2009). The decreasing cost of reading and writing microbial genomes has made it possible to generate billions of combinatorial genomic variants per day at specific loci (Wang et al. 2009; Isaacs et al. 2011; Jiang et al. 2013) and to sequence entire *E. coli* genomes for less than 25 USD per sample (Baym et al. 2015; Shapland et al. 2015) (**Supplementary Note 1**).

Computational analysis is increasingly a bottleneck when mapping whole-genome data to phenotypes across many samples. Going from raw DNA sequencing reads to annotated variants requires the integration of a large number of disparate tools, usually assembled into an *ad hoc* pipeline by individual labs and followed by time-intensive manual confirmation of variants. There remains a critical need for an integrated solution capable of comparative analysis among multiple genomes and supporting interactive querying and data visualization, collaboration, genome versioning, and the design of additional mutations or reversions (**Supplementary Table 1, Supplementary Note 2**).

To address this need, we developed Millstone, a web-based software platform that supports an iterative process of multiplex mutation analysis and genome engineering. Millstone automates read alignment and variant calling using a hybrid reference-based and *de novo* assembly approach, then allows researchers to explore and compare mutations among genomic samples, and finally creates updated reference genomes and designs new genomic edits for subsequent rounds of experiments (**Fig. 1a**). Serving as both a genomics pipeline and a platform for exploring whole-genome sequencing data, Millstone provides a powerful user-friendly interface that allows researchers to investigate individual variants through interactive filtering and alignment visualization (**Fig. 1b**).

**Figure 1:**
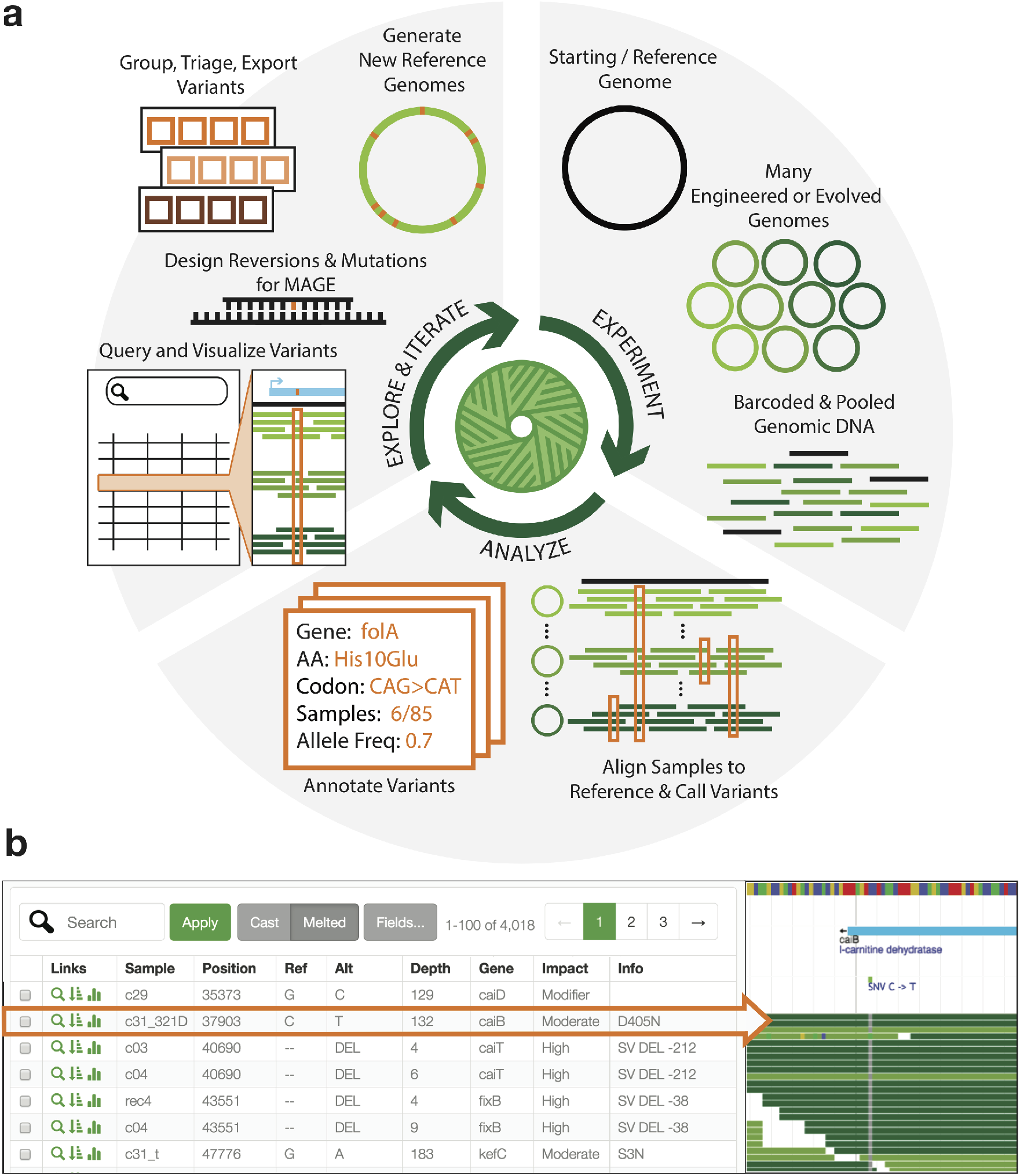
Millstone enables rapid iterative multiplex genome analysis and engineering. (**a**) To use Millstone, a researcher uploads a reference genome and next-generation sequencing reads for many individual genomic clones, for example from long-term evolution or targeted genome editing. Millstone performs alignment and variant calling for both single nucleotide variants and structural variation and then assigns predicted effects based on reference genome annotations. A unified data model stores sample genotype, phenotype, and variant annotation data. Variants can then be queried, filtered, and grouped into sets for export, triage, and analysis. These variant sets can be used to design oligonucleotides to recreate or revert mutations of interest, or used to generate new versions of the reference genome. (**b**) A combined screenshot of the Millstone analysis and alignment visualization views (condensed and cropped for clarity). A custom query language allows searching and filtering over the data. As variant calls sometimes require visual inspection and comparison, Millstone’s variant analysis view provides programmatically-generated links to visualizations of the relevant read alignments in JBrowse (Skinner et al. 2009).

## Results

Millstone was built in response to challenges encountered during the construction of the genomically recoded organism (GRO) *C*321. Δ*A* (Lajoie et al. 2013), a strain of *E. coli* in which all 321 UAG stop codons were replaced with a synonymous UAA. Multiplex automated genome engineering (Wang et al. 2009) (MAGE) was used to introduce sets of 10 mutations into 32 strains and conjugative assembly genome engineering (Isaacs et al. 2011) (CAGE) was used to hierarchically combine redesigned regions into a chromosome with all 321 UAGs recoded (**Fig. 2a**, green). We sequenced 68 intermediate clones to confirm the designed changes but our initial analyses were slow, error-prone, and lacked the ability to visualize and compare evidence for mutations among samples. Millstone solved these issues, allowing us to identify and track the 3127 designed and off-target mutations across all strains. Finally, by iteratively applying mutations directly to the initial reference genome and re-aligning reads, Millstone allowed us to generate a new *C*321.Δ*A* reference sequence which incorporated 355 additional off-target mutations that had accumulated during strain construction. (**Fig. 2a**, green and orange).

Millstone’s ability to rapidly generate clonal genotypes from whole genome sequencing reads enabled a follow-up project to improve the fitness of the GRO. The final strain from Lajoie et al. demonstrated incorporation of proteins containing non-standard amino acids, but suffered from an impaired growth phenotype, which we hypothesized was due to a subset of the 355 off-target mutations. We developed an iterative method for systematically optimizing strain fitness through predictive modeling and multiplex testing of reversions (*Kuznetsov et al., submitted*). Millstone was used throughout this process: first, to rank high-effect candidates for reversion, then to design oligonucleotides for MAGE, and finally to report variants from whole genome sequencing of 96 edited clones (**Fig. 2a**, orange). Once the final subset of effective reversions was identified and used to construct a faster-growing GRO, Millstone was also used to produce a final reference genome for the improved strain.

**Figure 2:**
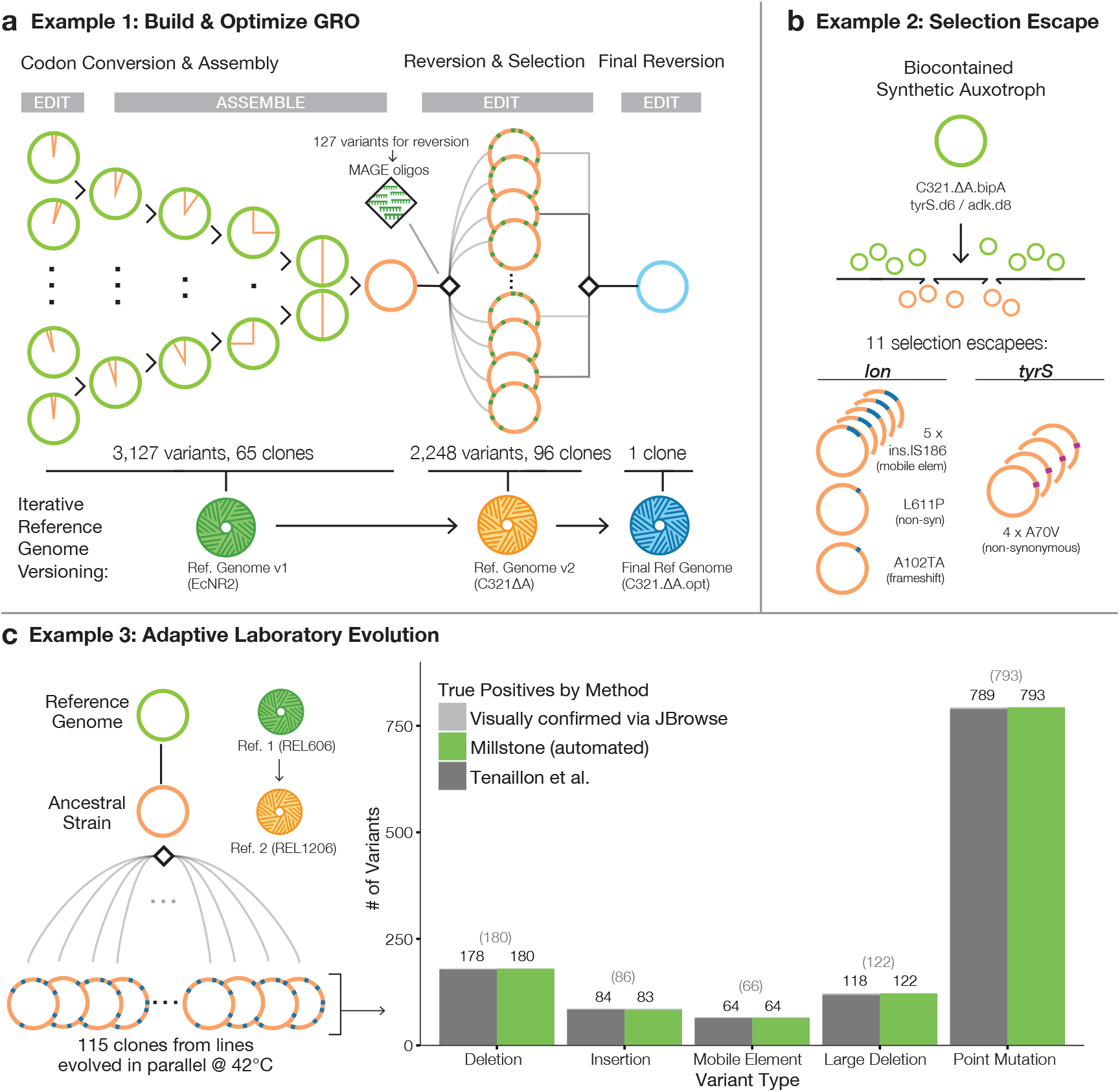
Millstone accurately detects genomic variants and can iteratively version genomes. (**a**) Millstone was used to analyze genomic clones involved in generating and rationally optimizing a genomically recoded organism. MAGE (Wang et al. 2009) and CAGE (Isaacs et al. 2011) were used to generate the *C*321.Δ*A* strain of *E. coli* (Lajoie et al. 2013). With sequencing data from these strains, Millstone confirmed the designed mutations, identified and annotated off-target mutations, and generated a new reference genome. Further reversion of variants was performed with MAGE to improve the strain’s fitness (*Kuznetsov et al., submitted*), and a final reference genome was generated. (**b**) Analysis of 11 escapee clones from a biocontainment selection with a synthetic non-standard amino acid (nsAA) auxotrophy (Mandell et al. 2015) identified two escape mechanisms, either mutation of *tyrS* or disruption of *lon*. (**c**) Millstone can also be used for Adaptive Laboratory Evolution studies. We employed Millstone to analyze mutations across 115 clones in the Tenaillon et al.(Tenaillon et al. 2012) high temperature evolution experiment. Millstone was used to create a new reference genome for the ancestral strain from REL606, the closest available reference genome, and called variants against this new reference. Millstone reports 99.2% of SNVs, deletions, and mobile elements found by the Tenaillon pipeline, as well some not identified in the original study (**Supplementary Table 2**).

Millstone’s *de novo* assembly and genotype comparison features were crucial in a project to engineer a biocontained version of the GRO which is dependent on a non-standard amino acid (nsAA) for survival (Mandell et al. 2015). A major challenge in engineering biocontainment, and in selection more generally, is diagnosis of escape mechanisms. In *Mandell et al*., plating of early versions of the biocontained GRO on non-permissive media revealed individual clones that could survive without the essential nsAA. We performed whole genome sequencing on 11 escapee clones and several controls and used Millstone to identify loci enriched for mutations across escapees. This led to the discovery and validation of two primary mechanisms of escape: a single off-target nonsynonymous mutation in the redesigned *tyrS* gene occurring in 4/11 clones and disruption of the *lon* protease in the remaining 7 clones. Millstone revealed several modes of *lon* disruption: a frameshift (1/7), nonsynonymous substitution (1/7), and insertion of a mobile element upstream of the gene (5/7) (**Fig. 2b**). To identify and precisely map these mobile elements and other structural variants, Millstone combines a local *de novo* re-assembly approach with coverage-based deletion calling (**Supplementary Fig. 3**). Rapid analysis of escapee clones allowed us to identify and validate the key mechanisms of escape from biocontainment, so that further modifications lowered escape rates by at least 5 orders of magnitude (undetectable escape with detection limit 2.2e-12 escapees per c.f.u.).(Mandell et al. 2015).

Millstone can also be used to analyze genomic variation in samples undergoing adaptive laboratory evolution (ALE). In Tenaillon *et al*. (Tenaillon et al. 2012), 115 lines of *E. coli* were grown at 42*^°^*C in parallel for over 2,000 generations in an attempt to identify convergent evolutionary responses to this environmental challenge (**Fig. 2c**). This impressive effort required a custom sequencing analysis pipeline consisting of over half a dozen tools, followed by manual validation and visual confirmation of all 1331 variants. We reanalyzed the raw data from this project in Millstone and identified 99.7% of SNVs and 98.9% of structural variants and mobile element insertions. Millstone further discovered 8 SNVs, 4 large deletions, and 2 mobile element insertions that were not reported in the original work (**Fig. 2d, Supplementary Table 2**). On an Amazon Web Services EC2 instance, the entire process from sample upload to variant triage across all 115 strains took a single day (**Supplementary Table 3**).

## Discussion

New technologies for constructing, screening, and selecting microbial genomes now allow for increasingly complex functional genomics studies and bioengineering endeavors. As the sequence constraints of the genome come into focus, the promise of designing new organisms to address medical and material challenges is becoming a reality (Haimovich, Muir, and Isaacs 2015). The path forward requires rapid construction and characterization of successive versions of redesigned genomes (Hutchison et al. 2016; Ostrov et al. 2016), and computational genome design and analysis tools will increasingly become integral to this process. Researchers who already have raw sequencing data can use Millstone to identify and explore mutations. We have reduced the barrier for other labs to use Millstone by making the software deployable on Amazon Web Services (AWS). Documentation and an online demo are available at http://churchlab.github.io/millstone.

## Methods

### Software availability and documentation

Millstone source code is available at https://github.com/churchlab/millstone. Documentation is available at http://millstone.readthedocs.io/.

### Deployment via Amazon Web Services (AWS)

Using Millstone via AWS is the recommended option for most users. We have pre-configured a Millstone installation as an Amazon Machine Image (AMI), allowing users to sidestep all dependency installation and configuration steps. Researchers can provision a fully-configured private instance of Millstone running on AWS in minutes and can specify compute, memory, and disk requirements to match project needs. AWS allows resizing a machine without losing data. For most projects and benchmarking, we use a 32-core r3.8xlarge instance type for alignment and variant calling, and then resize the machine to the 2-core c3.large or c4.large instance type for data exploration. Academic labs can apply for the AWS Cloud Credits for Research program (https://aws.amazon.com/grants/).

### Analysis pipeline

Millstone provides a user interface that guides a researcher through uploading a reference genome (Genbank or FASTA) and Illumina sequencing reads (FASTQ). The Millstone analysis pipeline then performs alignment, variant-calling, and annotation of called variants. Millstone aligns reads using BWA-MEM (Li and Durbin 2010), parallelizing across available cores. Millstone calls single nucleotide variants (SNV) using Freebayes and structural variants (SV) using a custom contig assembly-based method (see De Novo Assembly Pipeline below) and a custom coverage-based method. A diagram of the pipeline and the parallelization scheme is shown in **Supplemental Fig. 1**. Even though Millstone was primarily designed for haploid bacterial genomes, the default and recommended mode for calling SNVs in Millstone is diploid. This allows for marginal calls that are neither clean wild-type nor clean mutant alleles, indicated as “heterozygous” under the GT TYPE field (GT in VCF format). If the user-provided reference genome is annotated (i.e. Genbank format), the Millstone analysis pipeline can annotate variants with predicted effect using SnpEff (Cingolani et al. 2012).

### Variant exploration view

Millstone provides an interactive user interface for exploring mutations and comparing genomes. Two primary data view modes, *cast* and *melted*, allow exploring data with samples aggregated by mutation or by individual mutation-sample relationships, respectively. Each mutation row includes icons that link to the relevant view of aligned reads in JBrowse. This is useful for performing visual quality control (QC) on aligned regions to verify that reads are properly aligned around a variant or to diagnose complex structural events. Visual QC is particularly important for inspecting marginal variant calls (see Analysis Pipeline), where only a fraction of aligned reads show a SNV. These can indicate regional duplication, non-unique mapping of reads, or non-clonality.

A query language allows filtering variants according to fields such as read depth, evidence quality, gene affected, predicted impact (Online Documentation). The filter key GT TYPE is particularly important with respect to identifying mutant alleles and distinguishing them from marginal calls. GT TYPE can take on values of 0 (strong evidence wild-type allele), 1 (marginal call), or 2 (strong evidence mutant allele). The query language allows for boolean combinations of key-value statements. For example, the following query filters the displayed variants down to only well-supported mutants that have a moderate to strong affect on some gene:

~~~
GT TYPE = 2 & (INFO EFF IMPACT = HIGH | INFO EFF IMPACT = MODERATE)
~~~

Further information and examples of queries can be found in the Online Documentation (http://millstone.readthedocs.io/).

### Visualizing alignments

Millstone uses JBrowse (Skinner et al. 2009) to visualize read alignments, enabling manual quality control and verification of complex structural events. For each called variant, Millstone programatically generates a link that brings up JBrowse at the affected locus showing the relevant alignment tracks. In the *cast* view, multiple JBrowse tracks will be shown simultaneously if the variant is present in multiple samples.

### Variant sets

Variant sets are an important unit of operation in Millstone that allow grouping variants after filtering. Millstone’s analysis pipeline also allocates variants to several common variant sets by default, including sets indicating insufficient coverage, no coverage, greater than expected coverage, or poor mapping quality (corresponding perhaps to regions that are non-unique).

Variant sets can also be used to generate oligonucleotides targeting or reverting the mutations in the set, via an integrated python implementation of optMAGE (Wang et al. 2009) (https://github.com/churchlab/optmage). Millstone’s Genome Versioning feature allows for variant sets to be used to generate a new version of reference genome containing those variants as ‘ref’ alleles. In particular, we show the use of this feature to generate updated reference genomes for the C321.ΔA strain and its improved version, C321.ΔA.opt.

### De novo assembly pipeline

After Illumina sequencing reads are aligned to the genome, Millstone identifies candidate reads that may indicate structural variants, including unmapped, clipped, and split reads, as well as their pairs. These reads may indicate the presence of complex structural events such as deletions, novel sequence insertions, and translocations of mobile elements. Velvet (Zerbino and Birney 2008) is used to assemble these reads into *de novo* contigs.

Once the reads are assembled into contigs by Velvet, those contigs over a size threshold are aligned back to the reference genome using BWA-MEM (Li and Durbin 2010). These contig-to-reference alignments are used by Millstone to generate a graph whose nodes are contigs and reference sequence fragments and whose edges are alignments. Individual contigs and their reference junctions can be browsed and downloaded by the user. The contig sequences can also be downloaded as FASTA records and blasted individually by the user. Contigs whose edges map to annotated mobile elements are also labelled in the user interface. Novel sequence and mobile element insertions consist of multiple graph edges (two edges for novel sequence, four for mobile element insertions, **Supplemental Fig. 3b**). These variants are identified by a graph-traversal algorithm and converted into VCF records that are added to the variant database.

### Tenaillon et al. Variant Comparison

Raw data for Tenaillon et al. (Tenaillon et al. 2012) was downloaded from http://wfitch.bio.uci.edu/~tdlong/PapersRawData/Tenaillon.rawdata.tar.gz. Using the strain metadata provided with the sequencing data, we generated a targets file containing fitness ratios, line labels, and paths to read files for all 115 samples.

We first used Millstone to align the Ancestor strain (Line 0) to the REL606 reference genome (NCBI accession CP000819). This found a 6.93kb deletion in *scgB* (3,894,998 bp) and an IS186 insertion near *fimA* (4,517,603 bp). We used Millstone’s Reference Genome Versioning feature to generate the REL1206 reference genome by applying these strucutral events to REL606.

Finally, all 115 samples (including the Ancestor strain) were realigned to the new REL1206 genome using Millstone. The data were exported to CSV format for comparison with the variants called by Tenaillon et. al. For a list of mutation discrepancies between Millstone and Tenaillon, see **Supplementary Table 2**.

### Optimizing Genomically Recoded Organism C321

Δ**A** Millstone was used to create a new reference genome for the final C321.ΔA strain. Mutations called by Millstone were triaged based on quality and added to a variant set. Then, from the variant set view, the ‘Generate New Reference Genome’ feature was used to create the new genome. This process was iterated two additional times until there were no structural or single nucleotide variants called against the new final reference genome. 205

Millstone includes SnpEff (Cingolani et al. 2012) integration that allows annotating mutations with predicted effects. We used Millstone to generated 127 MAGE oligos to revert mutations predicted to have strong fitness effects.

After sequencing 96 clones which underwent from 5 to 50 cycles of MAGE with subsets of these 127 oligos, Millstone was used to align and call variants across all clones. This process took 3 hours on a 32 core machine (**Supplementary Table 3**). Millstone generated a .csv file reporting evidence for variants, and this was combined with per-strain doubling time measurements to generate a regularized linear model (Kuznetsov et al., submitted). Additionally, during modeling and iterative testing we returned to Millstone to use the JBrowse view to visually validate and compare marginal variant calls among samples.

Finally, Millstone was used to align, call variants, and produce a reference sequence for the final optimized strain, which contained 6 reverted alleles and 9 de novo mutations relative to the starting C321.ΔA strain.

### Mandell et al. Variant Comparison

We obtained sequencing reads from the authors and uploaded and aligned them to the C321.ΔA genome (described above). In addition to the 11 escapee genomes on the adk.d8 and tyrS.d6 strain backgrounds, we also sequenced control clones for each background which remained dependent on the non-standard amino acid bipA, and some additional escapee clones on other nsAA-dependent backgrounds.

To identify the SNVs, including the *lon* and *tyrS* A70V mutations, we used Millstone to look for non-designed mutations which occurred only in the escapee strains and not in the bipA-dependent strains. To exhaustively identify transposon insertions where only partial alignment support was available, we additionally downloaded the raw contig list from Millstone and looked for contigs with graph edges mapping from mobile elements to other genome locations. We identified junction contigs which bridge the IS186 mobile element and the *lon* gene in 5 strains (**Supplemental Fig. 3**).

### Data representation

Millstone stores variant, experiment sample, and evidence data across several tables in a PostgreSQL database (**Supplemental Fig. 2**). Returning results for user queries from the variant exploration view requires a ‘join’ operation among multiple tables, which is expensive and does not scale well. To address this, we use PostgreSQL’s Materialized View feature to compute and store a single denormalized table with all variant-sample evidence. Subsequent queries can then be performed directly against this table. Benchmarking on an AWS EC2 c3.large (2-core, 3.75gb RAM), a dataset with 100 samples and 2500 variants required 1 min 44 seconds to compute a materialized view and typical queries required less than 1 seconds. The Millstone database only recomputes the materialized view when underlying data has changed.

## Acknowledgements

We thanks members of the Church Lab for comments. Funding for this work was provided by U.S Department of Energy grant DE-FG02-02ER63445. G.K. and M.J.L. were supported by DOD NDSEG Fellowships. D.B.G. was supported by an NSF Graduate Research Fellowship. Computational resources for this work were provided by the AWS Cloud Credits for Research Program.

## Author Contributions

D.B.G., G.K., and M.J.L. conceived of and designed the software. G.K., D.B.G., B.W.A., K.Y.C., and C.C. wrote the software. D.B.G., G.K., M.J.L., and M.G.N. tested, debugged, designed individual features, and wrote documentation. D.B.G. and G.K. wrote the manuscript with input and edits from all authors. G.M.C. supervised the project.

## Supplementary Note 1: Cost of Multiplexed Genome Library Preparation and Sequencing

**Figure S1:**
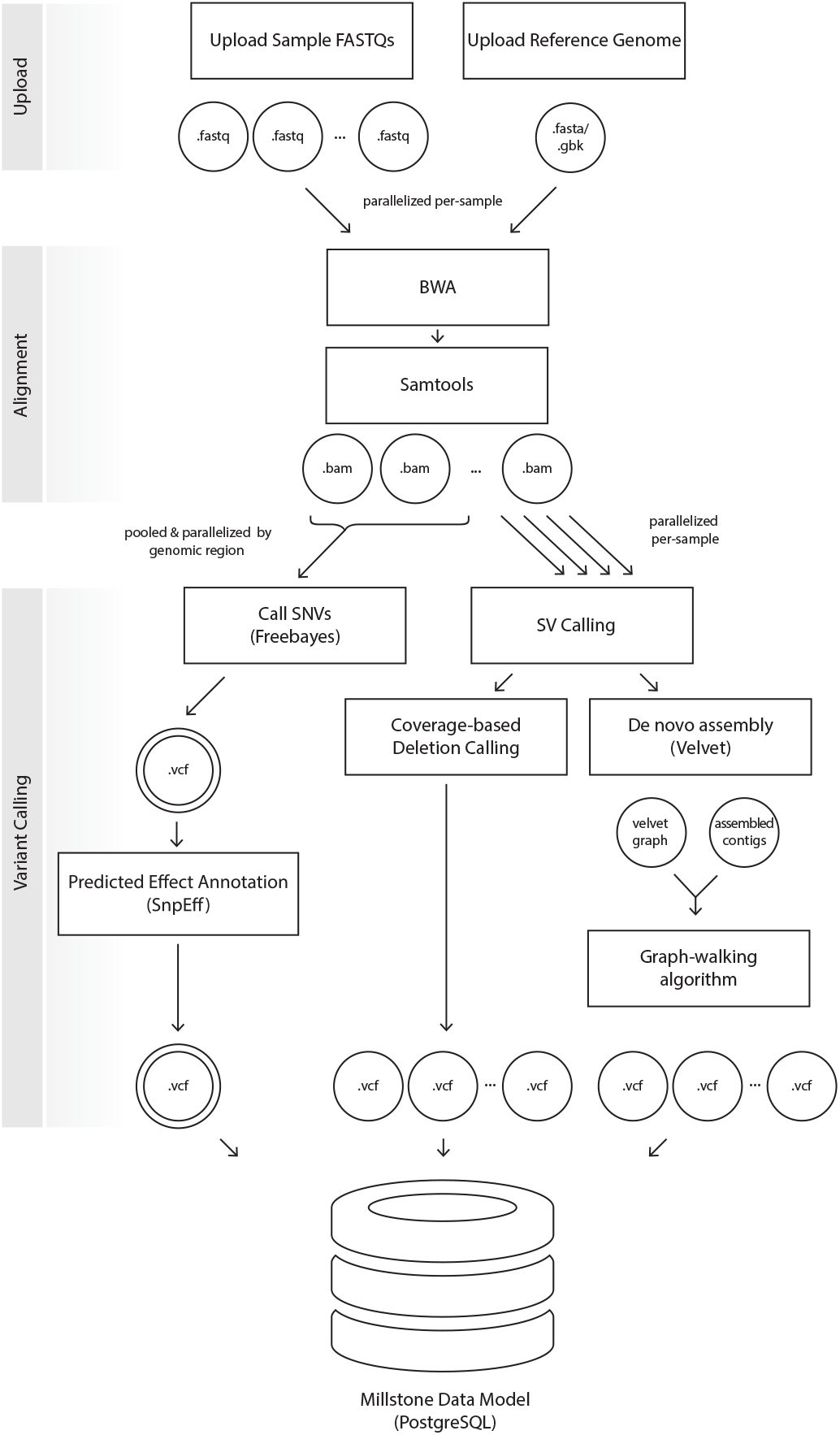
Millstone Analysis Pipeline. The analysis pipeline efficiently automates the process of identifying single nucleotide variants (SNVs) and structural variants (SVs) from user sample data and storing the information in a data model representation that can be explored using Millstone’s variant exploration UI. Once sample FASTQs and a reference genome are uploaded, Millstone uses BWA (Li and Durbin 2010) to align samples to the reference, parallelizing alignments across available processor cores. Once all alignments are complete, Freebayes performs SNV-detection on all .bam files simultaneously, parallelizing across regions of the genome. SVs are identified in parallel in individual samples using two methods: 1) Deletions are detected using sequencing read coverage and 2) novel junctions that indicate insertions and rearrangements are identified using *de novo* assembly of unmapped reads using Velvet (Zerbino and Birney 2008) followed by a custom graph-walking algorithm to combine assembled contigs and alignment with BWA to place contigs in the genome. All variant callers report their data in the Variant Call Format (VCF) and Millstone parses the VCFs into a single data model representation. Millstone further uses read coverage to identify regions of the genome with poor mapping quality and automatically adds variants that fall into such regions to appropriate VariantSets.

**Figure S2:**
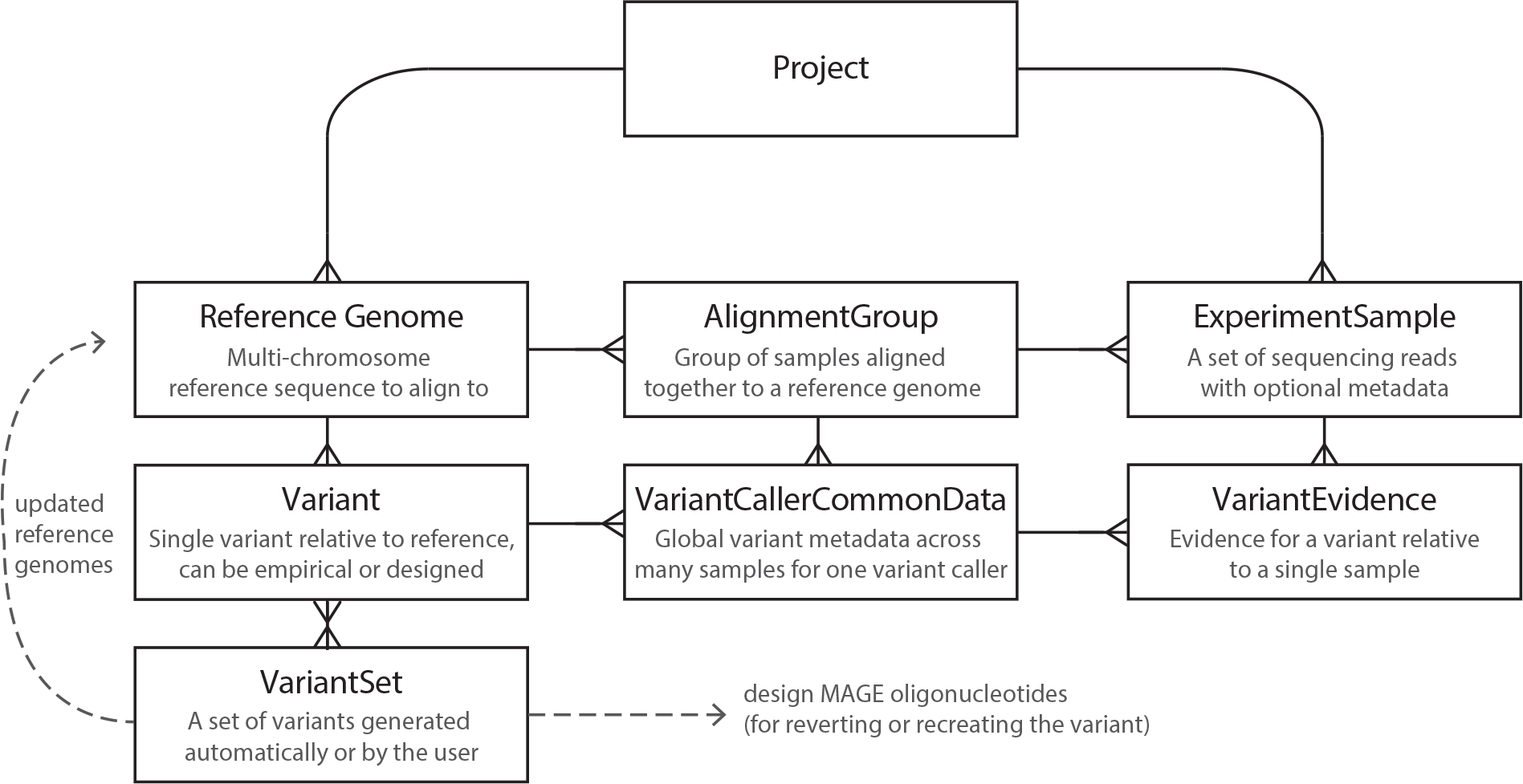
Millstone Data Model. Millstone’s data model was designed to enable project organization, data storage, and to support researcher operations including uploading data, running analysis pipelines, exploring the resulting data, and generating actionable outputs. Users upload *ReferenceGenomes* (e.g. Genbank or FASTA genome sequences) and *ExperimentSamples* (e.g. FASTQ) to a Project. The *AlignmentGroup* model stores data from an alignment of multiple *ExperimentSamples* against a specific *ReferenceGenome*. *Variants* represent both user-specified designed mutations and those emprically identified by variant callers. *Variants* are the most important primitive in Millstone, and serve as the unit of operation for analysis and design tasks. *Variants* are defined relative to a specific *ReferenceGenome*. The *VariantCallerCommonData* model relates a given Variant to any *AlignmentGroups* the mutation was called in and stores metadata provided by the variant calling tool (e.g. Freebayes). The *VariantEvidence* model further stores evidence for the occurrence of a specific Variant in a specific ExperimentSample. *VariantSets* allow the user to group Variants and take actions on groups. The VariantSet concept is very similar to tags in other software contexts and a Variant can belong to more than one VariantSet. Operations enabled by VariantSets include filtering in the exploration view, exporting subsets of variants, printing MAGE oligos, and generating new versions of reference genomes. The full data model is declared in the source code: https://github.com/churchlab/millstone/blob/master/genome_designer/main/models.py.

**Figure S3:**
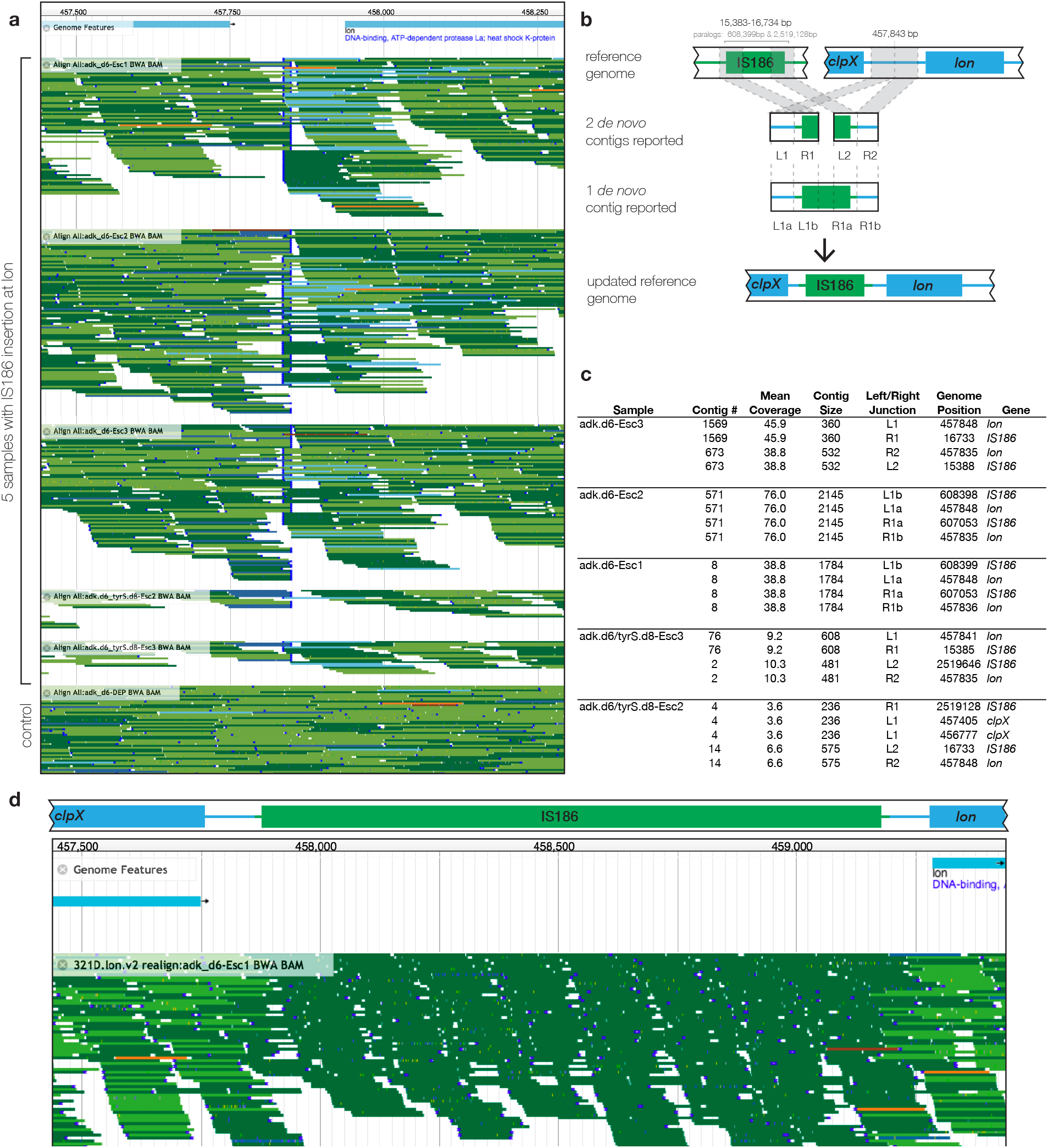
Millstone uses *de novo* assembly to identify a mobile element insertion at the *lon* locus across 5 escapee clones from Mandell *et al*. **(a)** Millstone’s integration with JBrowse shows evidence for a disruption upstream of the *lon* gene for 5 escapee strains. Each colored line represents a single read and the read alignments are grouped by sample. A wild-type strain is shown at the bottom for comparison. Darker reads map to the forward strand and lighter reads map to the reverse strand. Properly paired reads are green, reads with an unmapped mate are blue, and reads with improperly paired mates are orange. The dark blue vertical lines denote split reads, indicating a disrupted read alignment. **(b)** Millstone performs *de novo* assembly followed by alignment of assembled contigs back to the reference, and then uses a graph traversal algorithm to identify large insertions. Two example cases are shown where either one or two contigs are identified. These contigs are composed of reads that map to the *lon* locus and IS186 mobile element, indicating insertion of IS186 element at the *lon* locus. Finally, Millstone generates an updated reference genome reflecting the insertion. **(c)** A table of contigs, their sizes, and multiple reference alignments for each of the 5 samples with an IS186 insertion. **(d)** A new JBrowse view with the updated reference. The split and mismapping reads are gone, revealing a clean alignment in the region with the inserted IS element. The dark region indicates reads which map to multiple IS186 paralogs across the genome.

**Table S1:** **Comparison between Millstone and Other Tools**. A tabular comparison of features among different whole-genome sequencing tools, with a focus on those that are meant for use with haploid microbial genomes and are scalable to large datasets. All tools listed here are free and open-source. A more detailed description of the differences between the features and limitations of each is provided in **Supplementary Note 2**. *Effect Prediction*: Prediction of variant effects based on genome annotation. *Variant visualization*: can visualize read alignments for individual variants built into the tool. *Multiple Sample Comparison*: can compare the evidence for and presence/absence of a variant across multiple samples within the tool. *Interactive Querying*: can interactively search and subset variant list based on genomic position, gene, quality, mutation type, etc. within the tool. *Structural Variant Detection*: Supports detection of longer variants not normally detected by SNV callers like Freebayes and GATK Unified Genotyper, such as insertions, deletions, and translocations longer than 50-100 bp in length. *Genome Versioning*: Capable of generating an updated reference genome based on a subset of variants found in an initial reference genome. *Easy Deployment/Install*: Can be used without command-line compilation or scripting. *Genome Editing*: Generates designs for iterative editing/reversion of selected variants. *Sharing/Collaboration*: Built-in data-sharing via the web among teams of multiple users. *Supports Paired-End Data*: Utilizes paired-end read information to generate alignments and identify structural variants. *Modular Pipeline*: Capable of custom pipelines using different user-specified modules.

| Feature | Millstone | BreSeq | SPANDx | Galaxy |
| --- | --- | --- | --- | --- |
| Free & Open Source | • | • | • | • |
| Effect Prediction | • | • | • | • |
| Variant Visualization | • | • |  |  |
| Multiple Sample Comparison | • |  | • |  |
| Interactive Querying | • |  |  |  |
| Structural Variant Detection | • | • | • |  |
| Genome Versioning | • |  |  |  |
| Easy Deployment / Install | • | • |  | • |
| Genome Editing | • |  |  |  |
| Sharing / Collaboration | • | • |  | • |
| Supports Paired-End Data | • |  | • | • |
| Modular Pipeline |  |  |  | • |

**Table S2:**
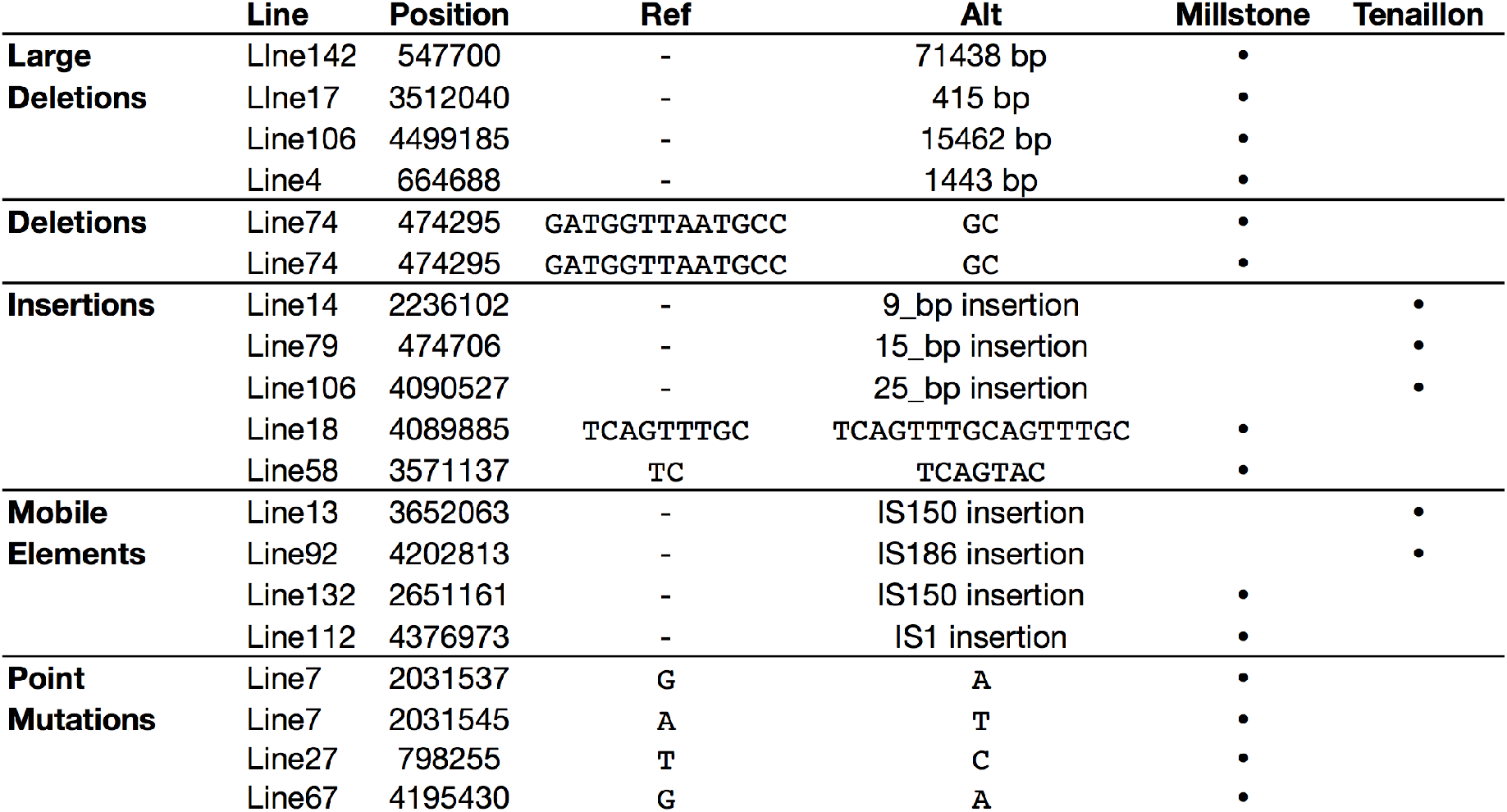
**Variant differences between Millstone and the original analysis performed by Tenaillon et al**. We compared variants found by *Tenaillon et al*. to those found automatically Millstone, focusing on Type II errors. Here we split discrepancies into 5 classes, including 3 short nucleotide variant (SNV) classes - short Deletions, Insertions, and Point Mutations, and 2 structural variant classes - Large Deletions and Mobile Element insertions. The final two columns describe ‘True Positive’ variants which were found by only one of the two pipelines. To identify these, we examined the read evidence using Millstone’s JBrowse visualization feature and determined whether the variant was correct as called by either pipeline.

**Table S3:**
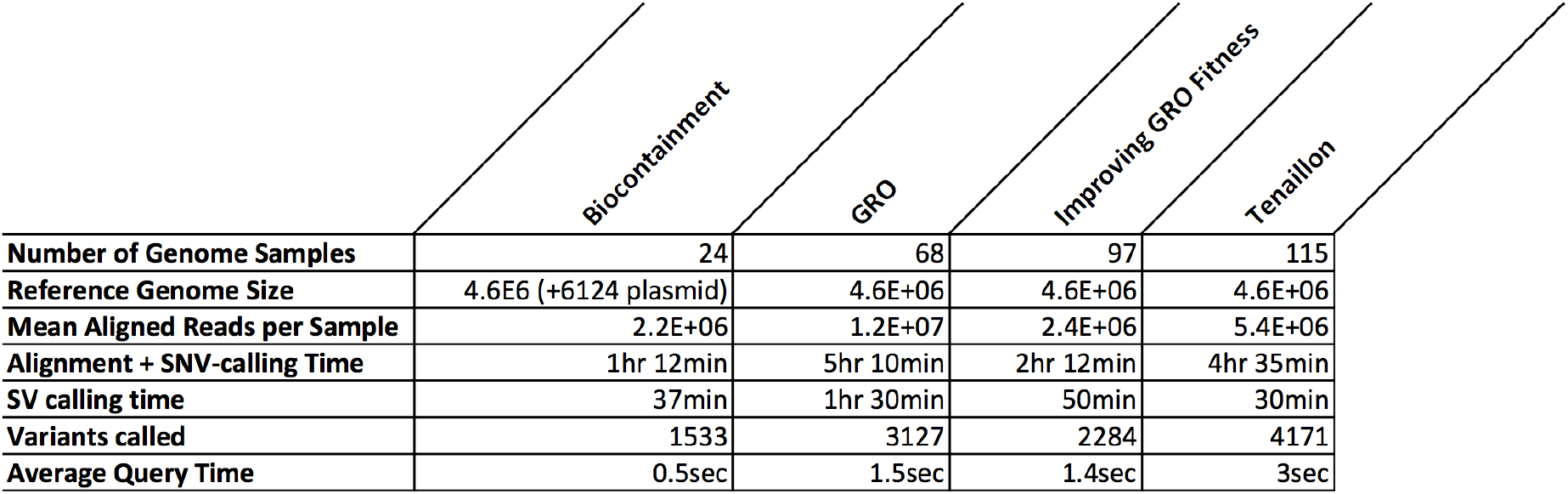
**Benchmarking**. Millstone’s analysis pipeline was executed on datasets of various size from four different projects: Biocontainment (Mandell et al. 2015), GRO (Lajoie et al. 2013), Improving GRO Fitness (*Kuznetsov et al., submitted*), and Tenaillon (Tenaillon et al. 2012). Average query time was calculated using no filter and a simple filter for strong alt calls: `GT TYPE = 2`. All benchmarking was performed on Amazon Web Services (AWS) Elastic Cloud Compute (EC2) instances r3.8xlarge (32 cores, 244 Gb memory).

|  | Biocontainment | GRO | Improving GRO Fitness | Tenaillon |
| --- | --- | --- | --- | --- |
| **Number of Genome Samples** | 24 | 68 | 97 | 115 |
| **Reference Genome Size** | 4.6E6 (+6124 plasmid) | 4.6E+06 | 4.6E+06 | 4.6E+06 |
| **Mean Aligned Reads per Sample** | 2.2E+06 | 1.2E+07 | 2.4E+06 | 5.4E+06 |
| **Alignment + SNV-calling Time** | 1hr 12min | 5hr 10min | 2hr 12min | 4hr 35min |
| **SV calling time** | 37min | 1hr 30min | 50min | 30min |
| **Variants called** | 1533 | 3127 | 2284 | 4171 |
| **Average Query Time** | 0.5sec | 1.5sec | 1.4sec | 3sec |

### Sample Preparation

Low-cost high-throughput sample preparation workflows for Illumina sequencing based on transposon insertion and fluorescent dye-based sample quantification can reduce the cost of preparation to below $15 USD per sample and be performed in approximately 5 hours per 96-well plate (Baym et al. 2015).

### Multiplexed Illumina Sequencing

For accurate discovery of structural variants, 20-30x coverage is ideal per sample. For the 4.6 megabase *Escherichia coli K12 MG1655* genome at 30x coverage, this is approximately 1 million 150 bp reads. The cost per read for Illumina sequencing can vary widely depending on the platform (NexSeq, MiSeq, HiSeq, etc). As a conservative estimate, a whole HiSeq 2500 lane in Rapid Run mode can produce 250 million paired end reads of 150 base pairs for a cost of approximately $2500 USD, or approximately $10 dollars per bacterial genome. Paired-end 150bp sequencing in Rapid Run mode takes approximately 40 hours. Larger-scale sequencing formats are generally cheaper, but longer to run, and smaller formats, like the MiSeq, are faster, but more expensive per genome.

## Supplementary Note 2: Comparison of Millstone to Other Tools

While other packages exist to solve the integration and automation of whole genome resequencing and annotation, most of these tools are built for large diploid genomes, such as *Homo sapiens*. Here we compare features and performance between Millstone and a few other related automated genome re-sequencing tools (see also **Supplementary Table 1**).

### Galaxy

Some tools, like Galaxy, allow users to create their own custom pipelines without bash scripting, and do support the creation of pipelines for microbial genomes. However, Galaxy requires that the user to understand and optimize settings for each individual tool. Galaxy also does not allow visualization or interactive querying of the output, and cannot generate new reference genomes or use the output of one round of sequencing to inform the next round. Finally, because of Galaxy’s one-size-fits-all nature, optimizing pipeline performance (for example, via inline compression and piping of input and output streams) is challenging.

### SPANDx

Another recent tool, SPANDx, can also perform genome resequencing for multiple strains simultaneously, but its widespread use is limited because it can only be run on UNIX computing clusters using the venerable and closed-source commercial PBS job scheduling system. SPANDx has no user interface or interactive components, and so users are required to gather the data manually and run the pipeline using a command line interface. Because we could not readily locate a PBS system to test the pipeline on, we were not able to compare the output between SPANDx and Millstone.

### breseq

*breseq* is purpose-built to perform haploid genome resequencing, and has become a standard tool for Adaptive Laboratory Evolution experiments. (Deatherage and Barrick 2014). *breseq* reports SNV and SV events for single genomes and provides a visualization of raw read evidence for the event. An advantage of Millstone is its ability to perform variant calling on hundreds of genomes in parallel, facilitating analysis of mutation frequency across a population of clones. Millstone’s JBrowse integration, data model, and querying features allow researchers to interactively filter, view, and compare genome alignments and variant data (as in **Supplementary Fig. 3**).

Both *breseq* and Millstone use a default variant detection threshold that works well in most cases, and Millstone complements this with an interactive search feature that allows researchers to filter variants after the variant calling pipeline has been run according to characteristics including read depth, number of samples with the variant, or predicted variant effect.

Millstone further supports paired-end reads, allowing for enhanced detection of structural variation, whereas *breseq* treats paired-end reads as single reads. Millstone can further assemble and place *de novo* contigs onto existing reference genomes. Millstone can be used pre-configured on Amazon Web Services (AWS) and so does not require proficiency with UNIX nor the manual installation of dependencies. Millstone’s user interface also automates the process of copying potentially large amounts of whole-genome data to the remote server.

